# Estimating the fraction of variance of crystallized intelligence explained by cortical surface area in early adolescence

**DOI:** 10.64898/2026.05.16.725604

**Authors:** Howon Ryu, Chun Chieh Fan, Armin Schwartzman

**Author notes:** Corresponding author: Division of Biostatistics and Bioinformatics, The Herbert Wertheim School of Public Health and Human Longevity Science, 9500 Gilman Dr, La Jolla, CA 92093, United States.

## Abstract

The relationship between cortical morphology and intelligence during adolescence has been widely studied, with existing literature reporting varying degrees of association across different modeling approaches. This study provides a comprehensive comparison of model performance in investigating the association between crystallized intelligence and cortical surface area using data from 11,351 subjects in the Adolescent Brain Cognitive Development (ABCD) study. We evaluate ten widely used models ranging from linear regression to graph convolutional networks across three covariate adjustment formulations: full (no adjustment), partial (age and sex adjusted), and total surface area (TSA) partial (age, sex, and TSA adjusted). Using bootstrap resampling with 50 iterations, we estimate the fraction of variance explained (FVE) for each model. Our results suggest that more complex models do not lead to higher FVE, with LASSO having the highest FVE of 15.9% (full formulation), Ridge at 10.5% (partial formulation), and Principal Component Regression (PCR) with 102 PCs at 2.5% (TSA partial formulation). Our results also reveal that the relationship between cortical surface area and crystallized intelligence is predominantly driven by global factors age, sex, and TSA, rather than by localized cortical surface area.

## 1 Introduction

The relationship between human cortical morphology and intelligence has been studied in the context of cognitive development in adolescence. While the link between cortical thickness and intelligence has been widely established [34, 10, 9, 28], there is a smaller volume of literature investigating the association between human cortex area and intelligence in adolescence, with focus being on how the cortical surface area, along with cortical thickness and gyrification, is associated with intelligence [39, 47].

The distinction of crystallized intelligence (*g*_*c*_) and fluid intelligence (*g*_*f*_) is a widely recognized concept in the theory of intelligence [13]. Crystallized intelligence (*g*_*c*_) encompasses a person’s accumulated knowledge and skills acquired from learning, as opposed to fluid intelligence, which refers to a person’s capacity to apply reason and logic to solve problems. Crystallized intelligence is also thought to be related to verbal ability [38] and openness [41, 52], a personality trait that suggests cognitive flexibility [18].

Cognitive outcome prediction from cortical surface information ranges from regression models to machine learning methodologies such as random forests or support vector machines (SVM) [42, 43, 5]. Deep learning is another popular approach for cortical surface inputs [48, 3]. In convolutional neural network (CNN) applications, cortical surface neuroimaging data are projected either to a 2D/3D grid [29, 16], or to a spherical space [50, 49] to account for the non-Euclidean geometry of the cortical surface. Some recent works also include transformer layers in their architecture [17, 14] or graph neural network architecture, which takes graph structure inputs [31, 32].

On the topic of the relationship between intelligence and cortical surface, existing literature reports varying degrees of association [15, 40, 33]. Applications using deep learning involve CNN [27] for crystallized and fluid intelligence prediction from cortical surface morphology, and fluid intelligence prediction from graph convolutional neural network [45].

While there is a plethora of studies explaining the association between human intelligence and cortical surface morphology with different combinations of methods used and human intelligence outcomes, our work has unique contribution in that: a) there is a lack of studies investigating specifically the association between crystallized intelligence and cortical surface area, and b) there needs to be an all-encompassing comprehensive investigation comparing the performances of models of differing complexities on a single dataset.

In this work, we investigate the link between cortical surface area and crystallized intelligence by estimating the fraction of variance explained (FVE) of crystallized intelligence by the cortical surface area in adolescence using the Adolescent Brain Cognitive Development (ABCD) study [22, 12]. We present FVE estimates using different statistical (linear regression, regression with regularization), machine learning (random forest), and deep learning (graph convolutional network) models. We compare model performance by evaluating the mean of the test R-squared estimates from the 50 bootstrapping samples as our FVE measure, after fitting the models on the training samples.

In our comparison, we include ten different models, from linear regression to the GCN model, with increasing model complexities. We also introduce a progressive approach in the covariate adjustments for each of our models: full, partial, and total surface area (TSA) partial formulations, where the total surface area is defined as the sum of cortical surface area over 20,484 vertices. With the partial formulation, we eliminate the effect of global factors age and sex. With the TSA partial formulation, we eliminate the effect of TSA on top of age and sex. This progressive approach in covariate adjustment is to investigate the localized effects or the brain sub-regions on crystallized intelligence. Figure 1 summarizes the analysis workflow for this paper.

**Figure 1:**
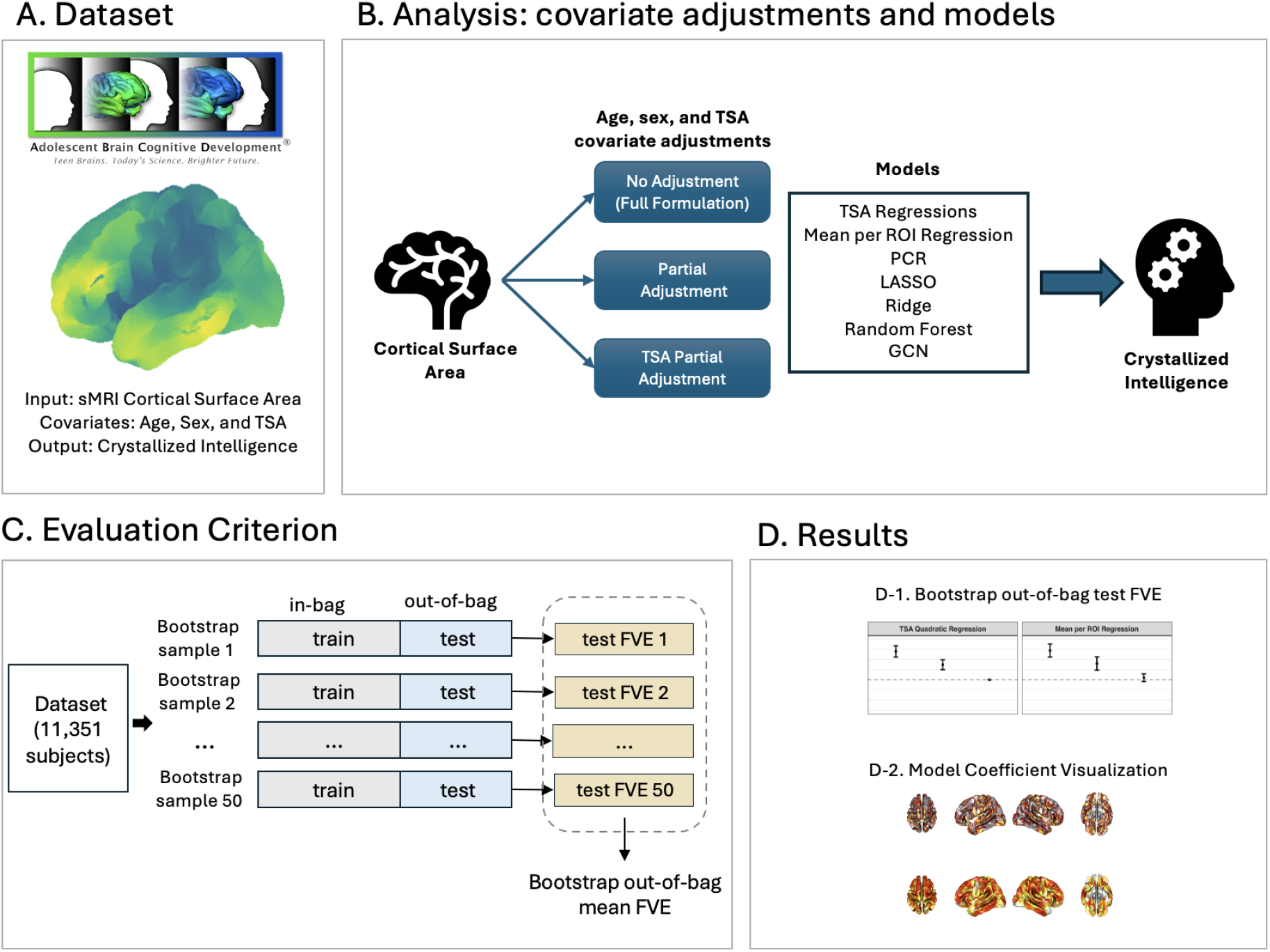
**A)** This analysis uses the Adolescent Brain Cognitive Development (ABCD) dataset with 20,484 cortical surface measurements from both the left and right hemispheres, crystallized intelligence scores, and age, sex, and total surface area (TSA) covariates from 11,351 subjects. **B)** Crystallized intelligence scores are predicted from the cortical surface area variables as defined in Table 1. The combinations of the three different covariate adjustments with ten models are investigated. **C)** Bootstrapping samples with replacement are drawn from the whole dataset containing the cortical surface information of 11,351 subjects. The model is trained on the in-bag samples, and the test FVE is calculated on the out-of-bag samples. After calculating 50 test FVEs from 50 bootstrap samples, the mean of the test FVEs are compared across models. **D)** Bootstrap out-of-bag test fraction of variance explained (FVE) is presented along with cortical surface mean coefficient plots as the model coefficient visualization.

This work reports the best FVE of 15.9% with no covariate adjustment, that is, including both age and sex as explanatory variables. After age and sex adjustment, the FVE is at 10.5%, while with age, sex, and TSA adjustment, the FVE drops substantially to 2.5%. Our work reveals more complex models do not increase the FVE, and that TSA along with age and sex are the major factors explaining the variability in crystallized intelligence.

**Table 1:** Summary of models and input cortical surface variables.

| Model | Cortical Surface Variables |
| --- | --- |
| TSA Linear Regression | TSA |
| TSA Quadratic Regression | TSA + TSA<sup>2</sup> |
| Mean area per ROI Regression | Mean area for each of 64 ROIs |
| Principal Component Regression (PCR) | Principal components |
| LASSO Regression | 20,484 cortical surface area values |
| Ridge Regression | 20,484 cortical surface area values |
| Random Forest | 20,484 cortical surface area values |
| Graph Convolutional Network (GCN) | Mean area for each of 64 ROIs |

## 2 Methods

### 2.1 Dataset

We take the demographic variables and neuroimaging information from the Adolescent Brain Cognitive Development (ABCD 5.1) dataset for young adolescents. The cortical surface area data from structural MRI (sMRI) are processed following the pipeline presented in [12, 22].

After additional data exclusion (excluding three records where sex is intersex-male), we base our analysis on 11,351 subjects. We use crystallized intelligence *g*_*c*_ measured by the NIH toolbox crystallized intelligence composite score [1] (range: 51-115; mean: 86.39) as our outcome variable, and the 20,484 cortical surface area values (range: 0-2.59 *mm*^2^) as our input variable along with age (range: 8.92-11.08, mean: 9.92) and sex (47.57% female) as covariates. Additionally, we consider the total surface area (TSA) which is a sum of cortical surface area over 20,484 vertices (left and right hemisphere combined) as another covariate.

Bootstrap sampling was used for repeated model fitting to produce FVE distribution for each model. The in-bag training dataset has size ⌊0.8 ·11, 351⌋ = 9, 080 with sampling with replacement from the 11,351 subjects. From the out-of-bag subjects, which are the subjects that were not chosen as the training dataset, test dataset is sampled with replacement with size ⌊0.2 · 11, 351⌋ = 2, 271.

### 2.2 Covariate Adjustments

The following section describes three types of covariate adjustment formulations (full formulation, partial formulation, and TSA partial formulation) which are repeated for each model presented in Section 2.3.

Let *Y* ∈ ℝ^*n×*1^represent the crystallized intelligence composite score (*g*_*c*_) outcome variable. Let *X* = {*X*_(*i,j*)_} ^*n×m*^ ∈ℝ^*n×m*^ be the design matrix with *m* cortical surface area variables, and *X*_*cov*_ ∈ ℝ^*n×*2^ be the design matrix containing the covariates age and sex. Together, *X*_*f*_ = [*X, X*_cov_] ∈ℝ^*n×*(*m*+2)^ represents the full design matrix containing *X*, and the covariates age and sex. The index for the subjects is *i* = 1, …, *n*, and the index for the cortical surface area variables is *j* = 1, …, *m*. Note here that *X* is a general term for different kinds of surface area variable, which is defined differently according to model type (See Section 2.3).

Then, let

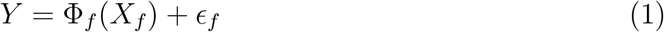

represent the ***full formulation*** that explains the variability in *Y* using the cortical surface area values and covariates with a differentiable function Φ_*f*_ (·), which is a general term denoting the models presented in Section 2.3.

There is a need to account for the multicollinearity between the covariates (age and sex) and the cortical surface area, where the relationship between the total surface area (TSA) and the covariates is known to be correlational [44, 26, 44]. For this, we perform a covariate adjustment formulation where the pure effect of cortical surface area on crystallized intelligence after accounting for age and sex is investigated. This is achieved by adopting the residuals 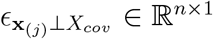 defined by the following linear regression models

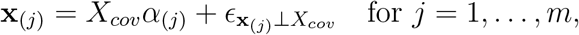

as the new independent variable. Here, the vector **x**_(*j*)_ ∈ ℝ ^*n×*1^ is the *j*^*th*^ column of the matrix *X*, and *α*_(*j*)_ ∈ ℝ ^2*×*1^ is the regression coefficient vector for **x**_(*j*)_ on *X*_*cov*_. In other words, 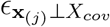are the residuals obtained from regressing (**x**_(*j*)_) on age and sex (*X*_*cov*_). These residuals 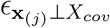 constitute the *j*^*th*^ column of the matrix 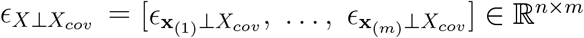 which becomes the new set of predictor variables in the ***partial formulation***

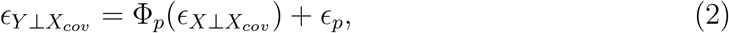

where Φ_*p*_(·) is a differentiable function, and the outcome *ϵ*_*Y ⊥X*_*cov* ∈ ℝ^*n×*1^ contains the residuals obtained from regressing *Y* on *X*_*cov*_ via the linear regression model

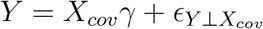

with the regression coefficient *γ* ∈ ℝ^2*×*1^.

Similarly, we further define an additional type of covariate adjustment formulation, the ***TSA partial formulation***, where we regress out the effect of total surface area (TSA) after accounting for age and sex. The TSA partial formulation is expressed as

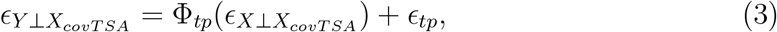

and is obtained in the same way as the partial formulation, with the only difference being *X*_*covTSA*_ ∈ ℝ^*n×*3^, the matrix containing age, sex, and TSA, instead of *X*_*cov*_.

Depending on the specifications of the model, the cortical surface area variable *X* can be one of the following:

a. TSA or TSA and TSA^2^, the quadratic value of total surface area
b. individual cortical surface area values with *m* = 20, 484 where the 20,484 columns represent each vertex
c. *m* number of principal components (PCs) from the principal component analysis (PCA) performed on the 20,484 area values
d. the mean cortical surface area value per 64 brain region of interests (ROIs) following the atlas presented in [24] which groups the vertices into 64 distinct areas including the background region.

Additionally, we also explore ***marginal baselines*** for linear regression to serve as a reference for the linear effect the covariates (age and sex) and TSA have on the crystallized intelligence. In the marginal formulations, we consider the following two models:

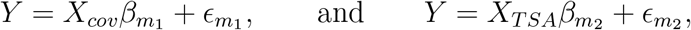

where *X*_*TSA*_ ∈ ℝ^*n×*1^ contains the TSA values, and 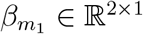 and 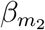 are the regression coefficients.

### 2.3 Models

Here, we introduce different models used in this analysis. All the models are formulated in three ways following formulations (1), (2) and (3) with the full formulation containing the covariates age and sex on top of the cortical surface variable. Model and the input cortical surface area variable pairs are summarized in Table 1.

#### 2.3.1 Baseline Models

**TSA Linear Regression** investigates the relationship between the outcome variable crystallized intelligence (*g*_*c*_) and total cortical surface area, which is the sum of the surface areas across all the cortical vertices. Additionally, **TSA Quadratic Regression** is performed with TSA^2^ term on top of the TSA variable.

**Mean area per ROI Regression** is another linear model with mean area per ROI as the input variables. According to the functional brain atlas suggested by [24] for the ABCD data, the 20,484 vertices are mapped into 64 distinct brain regions (64 ROIs plus background region). The resulting brain ROI map is presented in Figure 2. The mean cortical surface area values are calculated per each region, and linear regression is performed on the 64 mean area per ROIs as the independent variables, effectively reducing the input dimension from 20,484 to 64.

**Figure 2:**
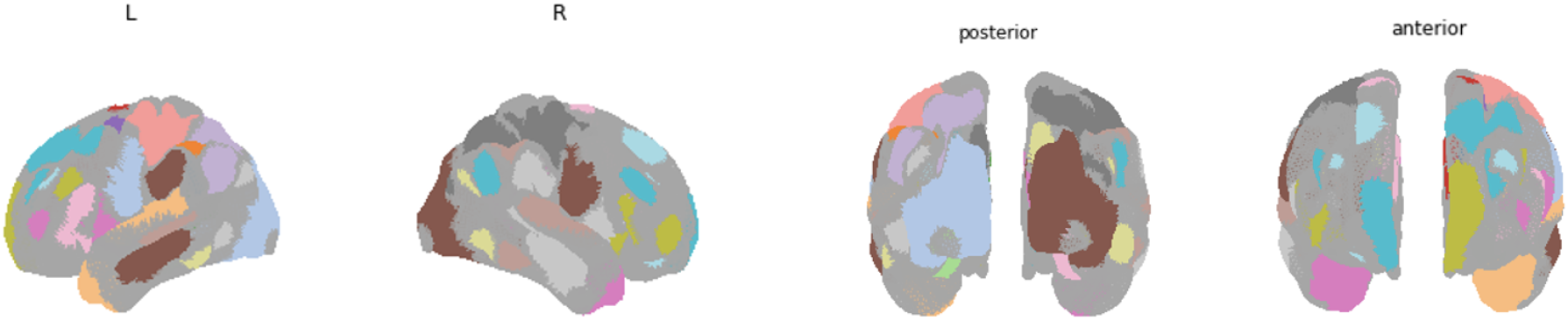
Cortical surface region of interest (ROI) mapping used for linear regression with mean area per ROI and GCN mean area per ROI. The mapping is adjusted from the functional brain atlas suggested by [24] to fit the 20,484 resolution and ABCD surface mesh ordering. There are 64 regions including 63 colored ROIs and the background (gray).

**Principal Component Regression (PCR)** is performed by first applying principal component analysis (PCA) on the standardized 20,484 cortical surface area values (and covariates for the full model) as dimension reduction. Then we consider taking 2, 102, and 1024 PCs, which represent 0.1%, 0.5%, and 5% of the initial dimension of 20,484, as predictor variables. Once the PCs are determined, linear regression is performed on the crystallized intelligence with the PCs as the predictor variables.

**LASSO (Least Absolute Shrinkage and Selection Operator) Regression** [53] is performed with outcome variable crystallized intelligence (*g*_*c*_) and the standardized 20,484 cortical surface area vertex variables along with sex and age covariates. The L1 regularization in the LASSO model effectively performs model selection, making it a good baseline in this high-dimensional setting. Five-fold cross-validation within the in-bag training set determines the optimized regularization parameter.

**Ridge Regression** [25], which is another regularized extension of linear regression, is performed with crystallized intelligence (*g*_*c*_) as the outcome variable and the standardized 20,484 cortical surface area vertex variables along with sex and age covariates. While Ridge regression does not shrink the coefficients to zero, the model mitigates the effects of multicollinearity by L2 regularization, making it another good baseline in this high-dimensional setting. Five-fold cross-validation within the in-bag training set determines the optimized regularization parameter.

**Random Forest** [8] is performed to capture complex non-linear relationship between crystallized intelligence (*g*_*c*_) and 20,484 cortical surface area variables. Random forest is a non-parametric learning method that constructs a large number of decision trees (100 trees in our case) which are aggregated to produce the mean prediction. Each tree is trained on a bootstrapped sample from the entire training dataset without sub-sampling, and all of the input features are considered at each split of a node. Node splitting was optimized via mean squared error.

#### 2.3.2 Graph Convolutional Network Model

To account for the structural and functional connectivity between the vertices and the spatial associations the surface area from each vertex has among themselves, we consider a Graph Neural Network (GNN) model.

Graph Neural Networks (GNN) are a class of deep learning models that takes graph-structured data as input. The graph relationship is represented by edges and nodes. GNN is well-suited when the non-Euclidean structure of brain surface should be accounted for, and is widely used in problems involving brain cortical surface morphology [20, 6, 46, 4]. We use a graph convolutional network model which uses a convolutional layer on the graph neural networks.

**Graph Convolutional Network (GCN)** uses the graph structure defined by the nodes, edges and the adjacency matrix. In this application, we use the mean area of the 64 ROIs as the inputs, with nodes representing the individual regions. The adjacency matrix is defined by the pairwise partial correlations between each pair of nodes (regions). The partial correlations are computed as follows. For any pair of regions *j* and *j*^*′*^, the cortical surface areas from all the other regions are linearly regressed out of the mean cortical surface areas **x**_(*j*)_ and 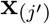, corresponding to the *j* and *j*^*′*^ regions respectively. Then, the Pearson correlation coefficient is computed between the two sets of residuals. This creates a fully connected graph.

Let 𝒢 = (*V, E*) represent a graph with *v*_*i*_ ∈ *V*, *i* = 1, …, 64, a set of nodes, and (*v*_*i*_, *v*_*j*_) ∈ *E*, a set of edges where edge (*v*_*i*_, *v*_*j*_) for *j* = 1, …, 64 is the connection between nodes *v*_*i*_ and *v*_*j*_. As stated above, the nodes are the 64 brain regions in our application. Then, *N*_*i*_ is the neighborhood of node *v*_*i*_ with *A* = {*A*_*ij*_}^64*×*64^ ∈ ℝ ^64*×*64^ representing the adjacency matrix defining the graph structure. The elements of *A*_*ij*_ are the edge features, defining the connectivity among the nodes.

In the GCN for any given subject, 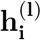 represents the embedded node feature vector for node *v*_*i*_ at the *l*^*th*^ layer. The first layer 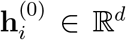 is the feature vector for node *v*_*i*_ which is the input for the model. In our application, 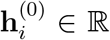 is the mean cortical surface area per region.

Across all the nodes, the embedded features can be expressed in the matrix form as 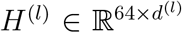, the *l*^*th*^ layer of the network. Following the formulation in [30], our GCN propagation can be expressed as

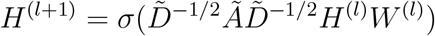

where 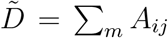, and *Ã* = *A* + *I*_64_ with *I*_64_ being the identity matrix of size 64. Learnable weight matrix is represented by 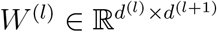 with the activation function *σ*.

Since the model goal of the GCN used is to predict *g*_*c*_, we incorporated the node pooling, read-out layer, and the regression head which is a fully-connected layer of dimension 1 for the final score prediction. The details of each layer can be found in [21, 19] for node pooling, and in [35] for the read-out layer.

### 2.4 Evaluation Criterion: bootstrap out-of-bag test FVE

We use bootstrap out-of-bag test FVE as our model evaluation criterion. Based on the whole dataset of *n* = 11, 351 subjects, 50 bootstrap samples are created by resampling 0.8*n* subjects with replacement. For each iteration of such bootstrap resampling, the in-bag 0.8*n* observations become the training set, and the 0.2*n* out-of-bag observations become the test set. The model is fit on the in-bag training set, and the FVE is calculated on the test set, yielding 50 test FVEs from each bootstrap sample. Mean and variance from the 50 test FVEs are compared across models.

### 2.5 Model coefficient Visualization

On top of the mean test FVEs from each model, we further investigate the model coefficients at the brain sub-region level from the four models with the highest FVEs (Mean per ROI Regression, LASSO, Ridge, and Random Forest). We consider the raw coefficients and standardized absolute coefficients. As the result of the model fitting process on the in-bag training datasets, Mean per ROI Regression produces 64 linear regression coefficients per bootstrap sample, LASSO and Ridge produce 20,484 regression coefficients respectively per bootstrap sample, and Random Forest produces 20,404 feature importance coefficients (range [0,1]) per bootstrap sample. The raw coefficients are defined as the average coefficient across the 50 bootstrap samples. Then, the standardized absolute coefficients are defined as the absolute value of the raw coefficients standardized to [0, 1] by min-max standardization.

We look into the distribution of the raw and standardized absolute coefficients and create cortical surface visualization, highlighting the patterns of the model coefficients per brain sub-regions. The visualization is performed using the standardized absolute coefficients, as it enables comparison across models despite the raw coefficients representing different mathematical measures.

We also show the mean (SD; standard deviation) Pearson correlation of model coefficients across models, which is obtained by correlating the raw coefficients on the vertex level across six pairs of different model combinations. Note here that since the raw coefficients from Mean per ROI Regression are defined on the ROI level, the coefficient values are divided by the number of vertices in the respective ROIs to represent the vertex-level coefficients.

## 3 Results

### 3.1 Bootstrap out-of-bag test FVE

Table 2 shows the out-of-bag test set mean and standard deviation of the FVEs from models with different covariate adjustments. Figure 3 summarizes Table 2 with mean and 2 standard error range visualization. Each model has full, partial, and TSA partial formulations to show gradual decrease in FVE as more covariates are accounted for. Models range from simple linear regression with TSA to GCN, increasing in model complexity.

**Table 2:** Comparison of model FVE across covariate adjustments: bold numbers are best performing FVEs in each covariate adjustment category.

| Model | Covariate Adjustment | Predictors | Mean FVE (SD) |
| --- | --- | --- | --- |
| Marginal Baseline |  | age+sex | 0.070 (0.008) |
|  |  | TSA | 0.055 (0.011) |
| TSA Linear Regression | full | TSA | 0.136 (0.014) |
|  | partial | TSA | 0.071 (0.012) |
|  | TSA partial | / | / |
| TSA Quadratic Regression | full | TSA + TSA<sup>2</sup> | 0.139 (0.014) |
|  | partial | TSA + TSA<sup>2</sup> | 0.074 (0.012) |
|  | TSA partial | / | / |
| Mean per ROI Regression | full | 64 ROIs | 0.143 (0.016) |
|  | partial | 64 ROIs | 0.080 (0.016) |
|  | TSA partial | 64 ROIs | 0.010 (0.009) |
| PCR (2 PCs) | full | PCs | 0.056 (0.011) |
|  | partial | PCs | 0.069 (0.013) |
|  | TSA partial | PCs | −0.001 (0.001) |
| PCR (102 PCs) | full | PCs | 0.087 (0.016) |
|  | partial | PCs | 0.096 (0.018) |
|  | TSA partial | PCs | **0.025** (0.014) |
| PCR (1024 PCs) | full | PCs | 0.061 (0.027) |
|  | partial | PCs | −0.011 (0.028) |
|  | TSA partial | PCs | −0.093 (0.030) |
| LASSO | full | 20,484 vertices | **0.159** (0.016) |
|  | partial | 20,484 vertices | 0.097 (0.016) |
|  | TSA partial | 20,484 vertices | −0.010 (0.021) |
| Ridge | full | 20,484 vertices | 0.124 (0.015) |
|  | partial | 20,484 vertices | **0.105** (0.017) |
|  | TSA partial | 20,484 vertices | 0.017 (0.017) |
| Random Forest | full | 20,484 vertices | 0.128 (0.009) |
|  | partial | 20,484 vertices | 0.092 (0.008) |
|  | TSA partial | 20,484 vertices | 0.020 (0.006) |
| GCN | full | 64 ROIs | 0.083 (0.058) |
|  | partial | 64 ROIs | 0.028 (0.038) |
|  | TSA partial | 64 ROIs | −0.009 (0.012) |

**Figure 3:**
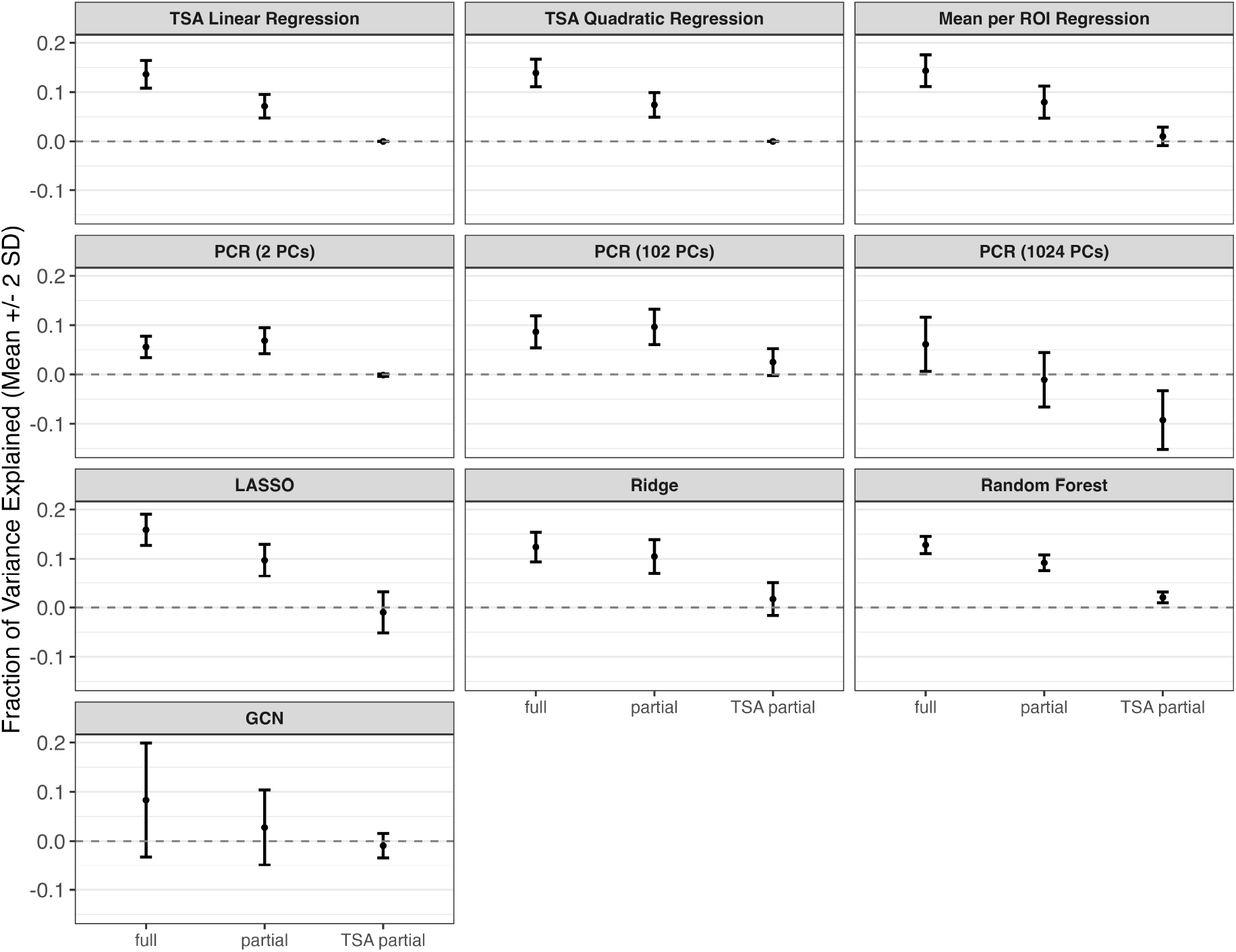
Mean FVE and 2 standard deviation range visualization for models with x-axis denoting different covariate adjustments.

The marginal effect of the age and sex is about 7.0%, whereas the marginal effect of TSA is around 5.5%. The FVE explained by the covariates and TSA is around 13.6%, as denoted in the TSA linear regression full model.

With no covariate adjustment (full formulation), LASSO regression achieved the highest mean FVE of 15.9%, followed by Mean per ROI Regression at 14.3%. After the age and sex covariate adjustment, Ridge regression showed the highest mean FVE of 10.5%, followed by LASSO at 9.7%. The TSA partial adjustment often results in negative FVE values, suggesting that the predicted *g*_*c*_ values from the model are often worse than when the model predicts the sample mean. With the TSA partial formulation, only the Mean per ROI Regression, PCR with 102 PCs and 205 PCs, Ridge, and Random Forest models showed FVE above 0%, at 1% to 2.5%.

As intended by the covariate adjustments, there is gradual FVE decrease from full to partial to TSA partial formulation within each model with an exception of PCR models. Overall, the decrease in FVE going from the full to the partial formulation is around 6-7% in TSA regressions, Mean per ROI Regression, LASSO, and GCN, which corresponds to the marginal effect of the covariates on *g*_*c*_. However, with Ridge and Random Forest the decrease is at 2-3%, showing high mean FVE at around 10% even with age and sex adjustment.

The decrease in FVE from the partial to TSA partial formulation is at another 7-8% for Mean per ROI Regression, PCR models, and Random Forest, whereas the drop is larger in LASSO and Ridge model at 11% and 9% respectively. GCN shows smaller drop at 4%. Since adjusting for TSA leads to trivial result for the TSA linear and quadratic regression (no variability in *g*_*c*_ explained by TSA after regressing out TSA), we leave the mean and standard deviation values as not applicable (/) for these models.

The PCR models show unstable FVE decrease from full to partial formulations, where PCR with 2, and 102 PCs show bigger FVE in the partial than the full formulation. It is also noteworthy that GCN, the most complex model in terms of number of trained coefficients, only shows 8.3% FVE with the largest standard deviation of all models (5.8%), suggesting the model coefficients varied highly depending on the training data. Overall, model complexity does not lead to higher mean FVE, as it is illustrated by the low mean FVE of GCN and high FVE of Mean per ROI Regression, despite having the same surface area predictor (mean area per ROI).

Based on the overall analysis result, the association between the crystallized intelligence (*g*_*c*_) and cortical surface area seems to be largely explained by the demographic covariates (age and sex) and TSA rather than localized effects of brain regions independent of TSA. This is especially evident when considering the fact that TSA partial models often show negative FVE values. Although models with more granular input surface variables show 2.0% (Random Forest), and 2.5% (PCR with 102 PCs) FVE for the TSA partial adjustments, they are still lower than the marginal effect of TSA (5.5%) or age and sex (7.0%).

### 3.2 Model Coefficient Visualization

#### 3.2.1 Model Coefficient Distributions

Figure 4 describes the distribution of the raw and standardized absolute coefficients from Mean per ROI Regression (denoted as”Mean per ROI”), LASSO, Ridge, and Random Forest. LASSO and Random Forest show the sparse and highly skewed standardized distribution, followed by Ridge and Mean per ROI regression.

**Figure 4:**
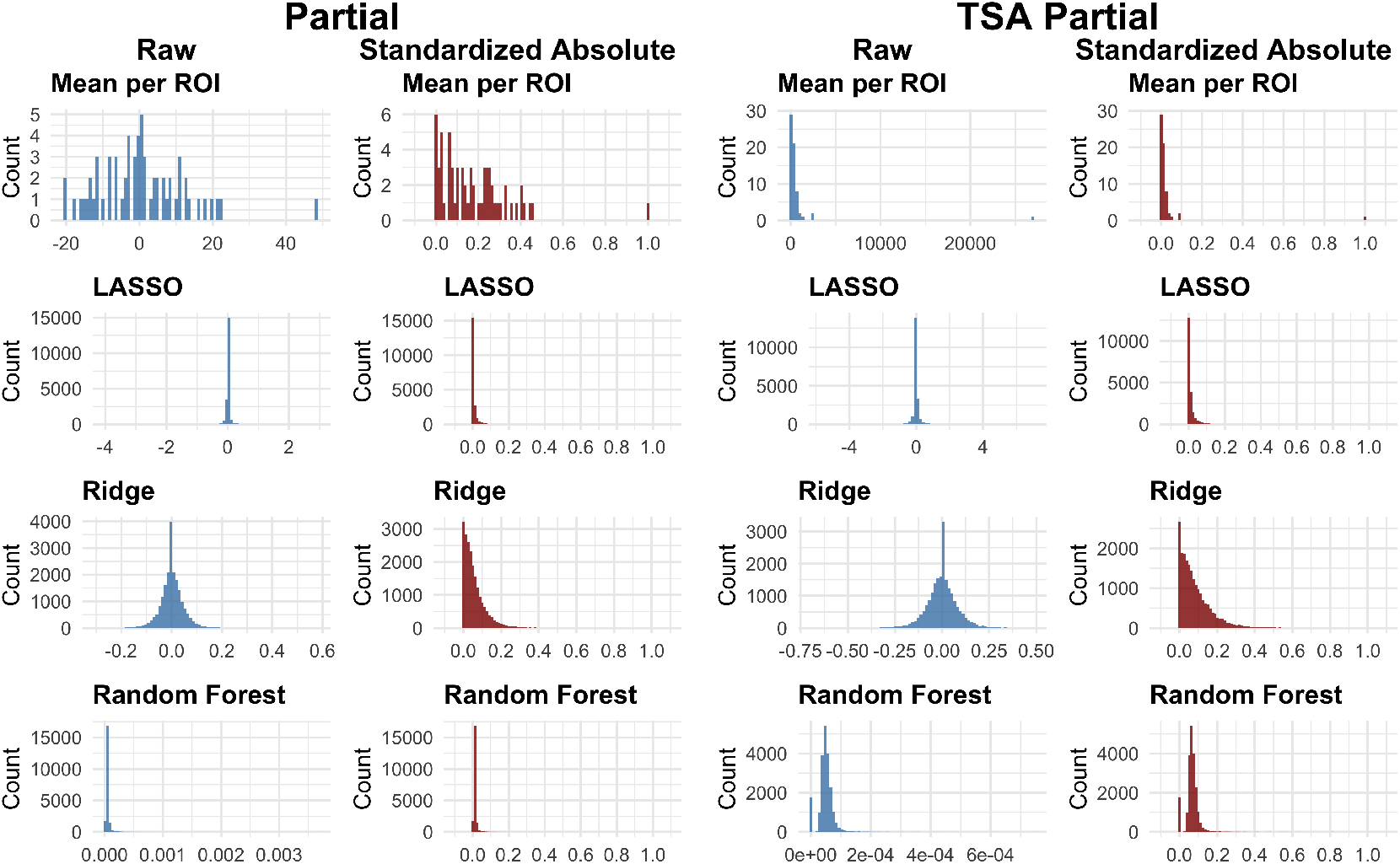
Histograms describing the distribution of the raw coefficients and the standardized absolute coefficients from Mean per ROI Regression (denoted as “Mean per ROI”), LASSO, Ridge, and Random Forest. The left panel shows the partial adjustment models, while the right panel shows the TSA partial adjustment models. The x-axis shows the coefficient value, and the y-axis shows the counts per coefficient value.

Mean per ROI Regression with partial adjustment shows relatively symmetrical raw coefficient distribution at 0 across the [-20, 20] range, with an exception of one ROI showing raw coefficient over 40. This translates into a skewed standardized absolute coefficient distribution. One notable thing is that in TSA partial adjustment, all the raw coefficients are positive, with an extreme outlier value above 20, 000.

In the LASSO and Ridge partial adjustment, the raw coefficients show normal distribution, with LASSO showing more sparse raw coefficients with more coefficients at 0 which is consistent with their L1 and L2 regularizations. This leads to a highly skewed standardized absolute coefficient distribution in lasso, with Ridge to a lesser degree. This pattern is repeated in the TSA partial adjustment, only with different range in the raw coefficient.

Random Forest raw coefficients in general show very small coefficients (*<* 0.001), again leading to very skewed distribution in the standardized distribution in both the partial and TSA partial adjustments.

#### 3.2.2 Model Coefficient Visualization on Cortical Surface

Figure 5 shows the cortical surface visualization of the model coefficients using standardized absolute coefficients. In Figure 5 left panel (partial), Mean per ROI Regression mostly highlights the background region as having the strongest standardized model coefficient, along with some ROIs showing mid-size effects. This pattern is replicated in Figure 5 right panel (TSA partial) with the background region highlighted as having a standardized absolute mean coefficient value close to 1. Across the two figures, LASSO has most coefficients suppressed to 0, while Ridge has smoother coefficients compared to LASSO, again in accordance with the characteristics of the L1 and L2 regularization. Random Forest shows model coefficient distributed across the whole brain.

**Figure 5:**
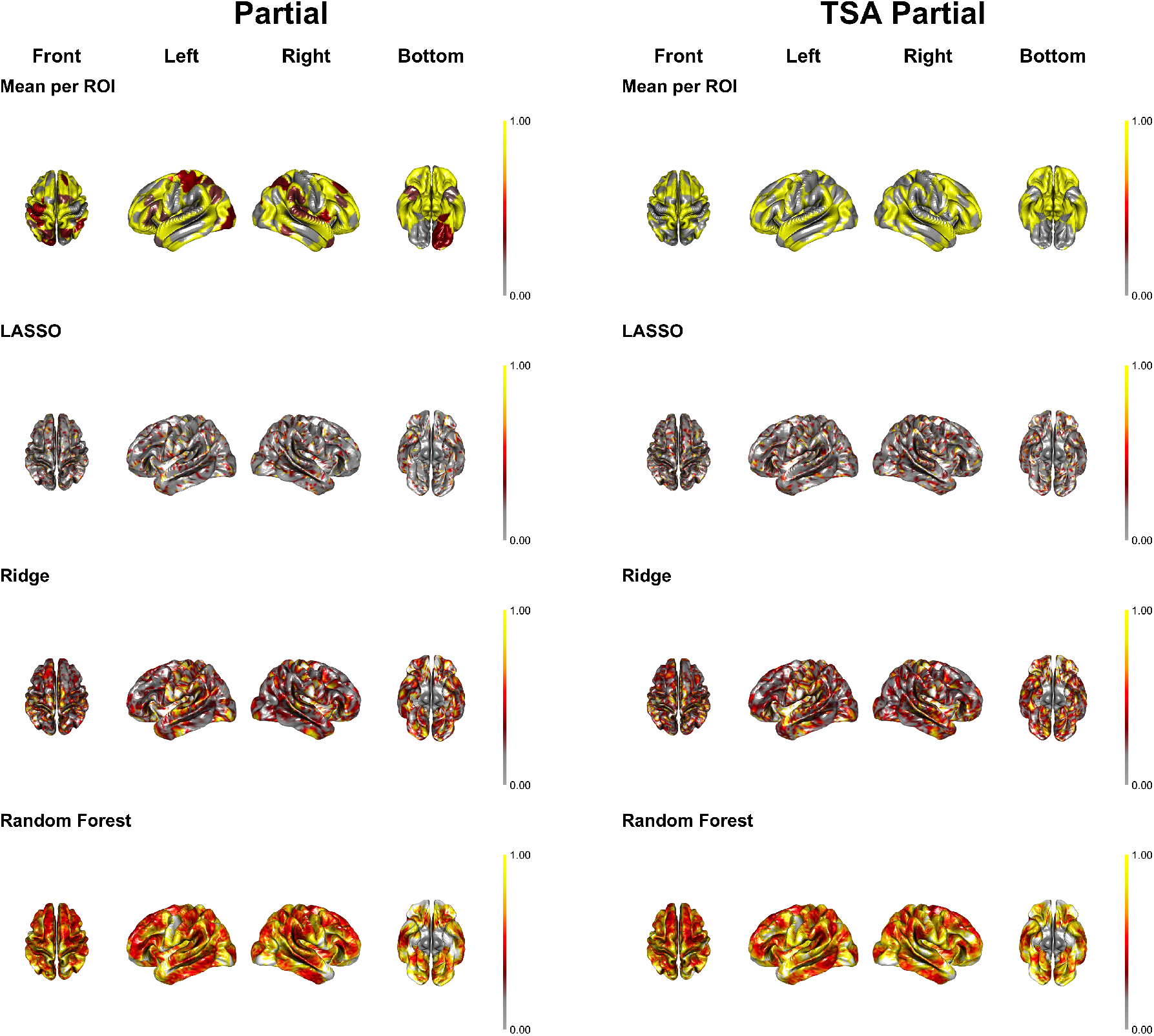
Cortical surface model coefficient plot for the partial (left) and TSA partial (right) adjustments: the absolute coefficients across 50 bootstrap samples are min-max standardized to [0, 1] range per model. Each column shows different view of the brain with each row representing the four models with the highest FVE.

Overall, no consistent pattern is observed across models as to whether a certain set of brain sub-regions is associated with the crystallized intelligence outcome. This again suggests that there is not enough evidence to conclude any sub-regional significance, but rather, the association seems to come from global factors such as age, sex, and TSA. Some existing literature shows similar conclusions consistent with our findings [51, 37].

#### 3.2.3 Model Coefficient Correlation between Models

To check how much the model coefficients of the brain sub-regions agree among the models, Figure 6 shows the mean (SD) model coefficient correlation over 50 bootstrap samples, and the correlation distribution.

**Figure 6:**
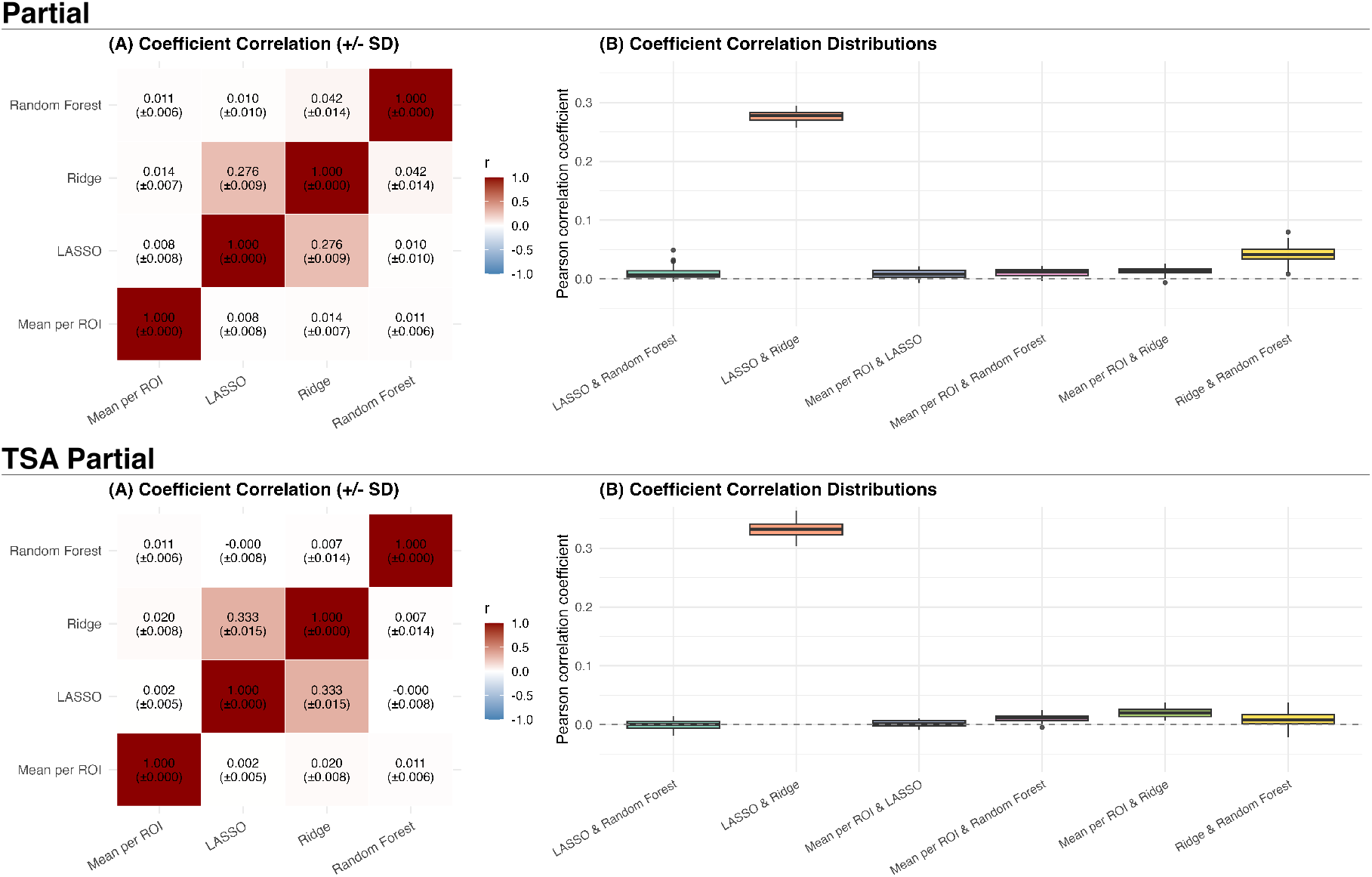
Model Coefficient correlation across models: the mean model model coefficient correlation (SD) across 50 bootstrap samples is displayed in the (A) plots, and the distribution box plot of the model model coefficient correlation is displayed in the (B) plots for partial (top) and TSA partial (bottom).

As prefaced by Figure 5, we fail to see overall high correlations among models, with only LASSO and Ridge pair showing the mean Pearson correlation coefficient over 0.1 at 0.276 (SD=0.009) for the partial adjustment, and at 0.333 (SD=0.015) for the TSA partial adjustment, as denoted in Figure 6 (A) panels. Figure 6 (B) panels show the distribution of the Pearson correlation coefficient over 50 bootstrap samples, showing overall low (*<* 0.1). The pattern of correlation coefficient among models is similar across partial and TSA partial adjustments.

This indicates that after accounting for global factors (age, sex, TSA), the estimated relationship between sub-regional cortical surface area and crystallized intelligence is inconsistent, with different models identifying different sub-regions as important rather than converging on biologically and neurologically meaningful brain regions. This again agrees with our conclusion that certain vertices or ROIs have large model coefficient due to model characteristics (such as regularization term), and not necessarily due to the inherent biological association.

## 4 Discussion

Our results do not show a prominent brain sub-region that comes up as important across models in explaining the crystallized intelligence. A contributing factor that makes direct comparison of models more nuanced is how different modeling approaches navigate the multicollinear features in different ways. With our input being the cortical surface area, we can assume a high level of spatial autocorrelation among the vertices due to the physical continuity of the cortical surface [36, 2, 11]. This explains the irregular coefficient patterns in LASSO and Ridge; in the presence of multicollinearity, LASSO’s L1 regularization tends to select one representative vertex from the surrounding correlated vertices and shrink others to zero, while Ridge’s L2 spreads coefficients more evenly across the correlated vertices. This does not occur in Random Forest or Mean per ROI Regression, as Random Forest aggregates information through multiple trees, allowing spatially adjacent vertices to contribute across different trees. Mean per ROI Regression avoids this problem entirely, as the inference is made on the pre-defined ROI level, wherein the vertices are assumed to be functionally correlated.

Our implementation of the GCN model used 64 regions as input rather than 20,484 vertices. We initially considered all 20,484 vertices as nodes, with the adjacency matrix defined as a binary matrix where the edge between two nodes was defined as 1 (connected) if the two nodes were connected in the 3D triangular mesh. However, this approach faced model fitting challenges. The high number of nodes (20,484 vertices) paired with low node feature dimension (dimension of one with cortical surface area) led to the model learning primarily from the node structure specified by the adjacency matrix, rather than node features. This ultimately led to low FVE due to the sparsity of the adjacency matrix defined by the physical brain structural connectivity in the 3D triangular mesh. We were able to improve fit by reducing the inputs to the 64 mean surface area values with the adjacency matrix defined by brain functional connectivity as suggested by [24].

In terms of FVE estimation, the final GCN estimated a lower FVE than the other less complex models even after incorporating both the structural (surface area node features from the sMRI) and functional (adjacency matrix defined by functional ROI map) features in our current GCN model. While the GCN methods are highly powerful in many applications, existing literature reports that graph neural networks do not always outperform simpler models in neuroimaging applications. [7] reports that when the spatial structure is prominent in the graph structure, Convolutional Neural Networks (CNNs) might outperform GNNs. [23] compares GNNs against classical machine learning methods on functional brain connectomes in neuroimaging studies, concluding that GNN models do not increase the predictive performance. In this sense, our result corroborates the claim that GNN models are not always ideal in neuroimaging applications, especially in the absence of well-defined graph structure as the input.

While the goal of our work was to present a comprehensive comparison of models in crystallized intelligence and cortical surface area association, some limitations warrant consideration for future work. First, our analysis focused exclusively on cortical surface area as the sole cortical feature at a single time point. Future work could investigate incorporating multiple cortical properties (thickness and gyrification), or using longitudinal design to model developmental trajectories. Additionally, model fitting and validation across multiple datasets spanning different populations would strengthen the generalizability of our modeling approach.

## 5 Conclusion

This study provides a comprehensive investigation into the fraction of variance of crystallized intelligence explained by the cortical surface area depending on different modeling methods. Bootstrapping mean test FVEs with three different covariate adjustment formulations were compared across ten models. Cortical surface mean coefficient plots were presented to show brain sub-region level effect size visualization.

LASSO achieved the highest FVE at 15.9% with no covariate adjustment (full formulation). After adjusting for age and sex (partial formulation), Ridge regression showed the highest FVE at 10.5%. Under the TSA partial adjustment, FVEs dropped substantially across all models, with PCR (102 PCs) achieving the highest at 2.5%.

The results suggest that the relationship between cortical surface area and crystallized intelligence, under our given sample size, vertex resolution, and single time point inputs, is predominantly driven by the global factors age, sex, and TSA, rather than by localized cortical surface area. TSA partial adjustment models frequently show near-zero to negative FVEs, indicating that little to no variance is attributed to factors beyond age, sex, and TSA. The cortical surface mean coefficient plots further corroborate this conclusion with no coherent sub-regional patterns in effect size.

## 6 Code Availability

The code used for simulations and data analysis can be found in the github repository.

## 7 Acknowledgements

This was partially by the National Institutes of Health [R01MH128923 to A.S. and H.R.] The ABCD data repository grows and changes over time. The ABCD data used in this report came from NIMH Data Archive Digital Object Identifier (DOI) 10.15154/z563-zd24. DOIs can be found at https://dx.doi.org/10.15154/z563-zd24.

